# Dynamical Systems Approach to Evolution-Development Congruence: Revisiting Haeckel’s Recapitulation Theory

**DOI:** 10.1101/2020.11.17.387506

**Authors:** Takahiro Kohsokabe, Kunihiko Kaneko

**Affiliations:** Laboratory for Evolutionary Morphology, RIKEN Center for Biosystems Dynamics Research (BDR), Kobe, Japan, 650-0047; Research Center for Complex Systems Biology, Graduate School of Arts and Sciences, The University of Tokyo, Tokyo, Japan, 153-8902

**Author notes:** Correspondence to: Kunihiko Kaneko, Research Center for Complex Systems Biology, Graduate School of Arts and Sciences, The University of Tokyo, Tokyo, Japan.

## Abstract

It is acknowledged that embryonic development has tendency to proceed from common toward specific. Ernst Haeckel raised the question of why that tendency prevailed through evolution, and the question remains unsolved. Here, we revisit Haeckel’s recapitulation theory, i.e., the parallelism between evolution and development through numerical evolution and dynamical systems theory. By using intracellular gene-expression dynamics with cell-to-cell interaction over spatially aligned cells to represent the developmental process, gene regulation networks (GRN) that govern these dynamics evolve under the selection pressure to achieve a prescribed spatial gene expression pattern. For most numerical evolutionary experiments, the evolutionary pattern changes over generations, as well as the developmental pattern changes governed by the evolved GRN exhibit remarkable similarity. Both pattern changes consisted of several epochs where stripes are formed in a short time, whereas for other temporal regimes, pattern hardly changes. In evolution, these quasi-stationary generations are needed to achieve relevant mutations, whereas in development, they are due to some gene expressions that vary slowly and control the pattern change. These successive epochal changes in development and evolution are represented as common bifurcations in dynamical systems theory, regulating working network structure from feedforward subnetwork to those containing feedback loops. The congruence is the correspondence between successive acquisition of subnetworks through evolution and changes in working subnetworks in development. Consistency of the theory with the segmentation gene-expression dynamics is discussed. Novel outlook on recapitulation and heterochrony are provided, testable experimentally by the transcriptome and network analysis.

## Introduction

Even before Darwin’s theory of evolution was generally accepted, a common trend in embryonic development across species had already received attention. von Baer (1828) observed that during embryogenesis, form of embryo diverges from a common shape shared across many species and eventually become species-specific. In other words, embryos of many species within the same phylum possess a common developmental stage, then undergo species-specific changes throughout the development. Haeckel studied development in the context of Darwin’s evolution theory. According to him, the similarities in embryonic development at the early stages stem from the common ancestor of species, whereas new stages towards the end of the developmental process are due to adaptation in evolution. He hypothesized that there is a tendency in evolution of developmental processes, that is, the early developmental stages are ancestral, hard to change and thus conserved, whereas the later stages are derived and easy for further alteration. Thus, he claimed, development is phyletically constrained to be parallel to evolution.

However, Haeckel was unable to validate his hypothesis using quantitative and comparative analyses as the role of genes as the origin of inheritance and the epigenetic regulation of development remained undiscovered during his time. Thus, quantitative examination of developmental and evolutionary processes was not possible.

Presently, developmental biology has made great advances, and developmental processes are now studied in terms of molecules and genetics. A century after Haeckel, quantitative comparative analysis of development across species is now available. Furthermore, it is uncovered that at a phylotypic stage, i.e., the middle phase of embryogenesis, the similarity among species of the same phylum is maximal (Hazkani-Covo, Wool and Graur, 2005; Irie and Sehara-Fujisawa, 2007; Domazet-Lošo and Tautz, 2010; Kalinka et al., 2010; Irie and Kuratani, 2011; Quint et al., 2012; Levin et al., 2012; Wang et al., 2013). The existence of the phylotypic stage elucidated the significance of the independence of each phylum and the conservation of body plans, which is a viewpoint different from classical morphology. After the phylotypic stage, the similarity decreases as species-specific developmental changes occur. The phylotypic stage is the bottleneck of developmental similarity, and embryogenesis has been often termed the “hourglass model” (Sander, 1983, Duboule, 1994, Raff, 1996). Although it is not identical with Haeckel’s theory, the evolutionary divergence of traits after the phylotypic stage may be regarded as a modified version of his theory.

In spite of the advances in the field of developmental and evolutionary biology over the past one hundred years, the general developmental tendency from the common toward the specific is not yet established. How the tendency, if it exists, is shaped by evolution remains to be resolved. One of the reasons for the difficulty to answer to this question is the poor data on the evolution of developmental processes. Paleontological records are usually sparse, and genomic/ embryonic information are often lost. Comparison of the development among currently-living species is possible. However, there is no guarantee that species that retained ancestral traits also retained the developmental process of their ancestors; the developmental process of current-living species could have changed via the adaptation to the environment (cf. Developmental Systems Drift, True and Haag, 2001). Thus it is hard to determine homology of the developmental process, particularly regarding the ancestral states.

Moreover, even if reasonable hypothesis could be proposed from these data, it is not possible to test if they hold true even when evolutionary tapes are rewound. We cannot experimentally set up the conditions of passed times; even if it was possible, it would take an enormous amount of time to study the evolution. Then, even if the complete recapitulation or any other evolutionary-developmental relationship would be observed from the data, it would be quite difficult to distinguish whether they happened by chance or due to a necessity of evolution.

Computer simulations of evolution can be a strong aid for this situation. Here, we first prepare a population of individuals of different genotypes. Genes determine the spatial pattern dynamics to shape the phenotype, via the gene expression dynamics of cells. The fitness of each individual is determined by a particular function of phenotypes, i.e., spatial patterns of given gene expression over cells. The genotypes that encode phenotypes of higher fitness are selected, and subject to mutations; slight changes in the rule of gene expression dynamics. Thus the population consists of slightly different genotypes, from which the fitness are computed again to select the next generation. With this setup, evolution can be mimicked. Of note, as the phenotype is shaped as a result of developmental dynamics, one can examine what types of developmental dynamics evolve over generations, under a given fitness constraint on the phenotype.

These evolutionary simulations are advantageous as literally complete data through evolution are available: one can keep the track of the pedigree as a chain of mothers-to-daughters (see Fig.1). Analysis on such single-chain-phylogeny gives further species-wide comparison, which is possible only in computer simulations.

**Figure 1.**
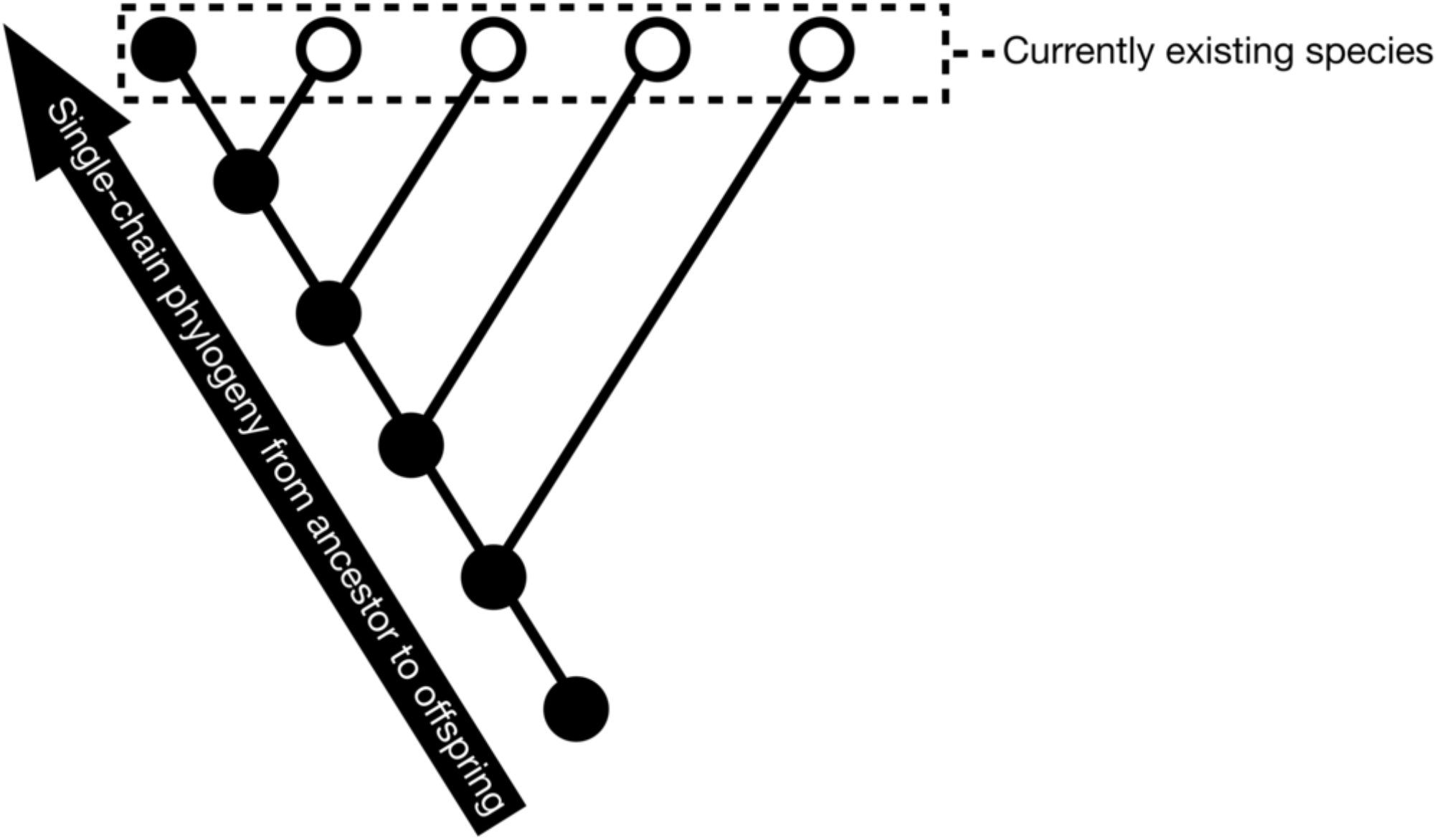
Schematic representation of the single-chain phylogeny. In the phylogenetic tree above, the currently existing species, which are denoted by the circles inside the dotted line, originate from a common ancestor at the root. The comparison of the developmental processes across species are usually carried out over these existing species. This study focused on the comparison along a single phylogenetic chain, which is represented by the black dots. The comparison of developmental processes along this chain is possible at least in theory or simulations, which provides a fundamental information on the possible relationship between evolution and development.

As another advantage, these computational simulations are repeatable so that with multiple simulation runs, we can examine the generality of a particular evolution-development relationship that is found in a single-run simulation, by repeating the runs later. Computer simulations can capture such mechanisms that are relevant to real biological systems and/or plausible evolutionary trails.

Simulation of gene regulation networks has first been studied at the single-cell level,(Glass and Kaufman, 1973; Mjolsness, Sharp and Reinitz, 1991). Evolution of gene networks has been carried at the single-cell level, where a certain gene expression pattern was assigned as a fitness. Extensive simulations have been carried out, to study the evolution of robustness to noise and to mutations (Wagner, 2005; Ciliberti, Martin and Wagner, 2007; Kaneko, 2007), evolution of certain network structures or modularity (Ma et al., 2009; Inoue and Kaneko, 2013), and so forth.

Computer simulations of developmental process also has been studied by including a spatial pattern consisting of multiple cells aligned on a one-dimensional space. For instance, the on/ off patterning of gene expressions on 1-dimensional space governs segmentation, as was reported (Salazar-Ciudad, Newman and Sole, 2001a b). Two mechanisms, i.e., feed-forward regulation from maternal gradient, as in Drosophila, and the oscillation-fixation mechanism have been elucidated (François, Hakim and Siggia, 2007; Fujimoto, Ishihara and Kaneko, 2008). Importantly, the mechanisms explain the simultaneous segmentation that is observed in development in long-germ cells as in Drosophila and the sequential segmentation as in vertebrates. The segmentation that precedes the speciation of body parts is also discussed (ten Tusscher and Hogeweg, 2011). The mechanisms elucidated in these simulations are relevant to the understanding of the segmentation processes in real biological systems as well as their evolutional origins (ten Tusscher, 2013).

In these studies, pattern formation dynamics result from gene expression changes through the evolution of gene regulation networks. Of note, using the protocol of these simulations, the developmental change in the gene expression pattern can be compared with the evolutionary change. This strategy of numerical evolution-development comparison opens us the possibility to examine the recapitulation in a quantitative and rigorous fashion. Below, we summarize the recent advances in recent advances in dynamical-systems theory and evolutionary simulations, with which recapitulation is explained as a result of control by slowly varying gene expression dynamics. Some experimental observations in segmentation evolution in arthropod are discussed accordingly, whereas experimental verification of the theory by transcriptome analysis is suggested. Furthermore, heterochrony is explained according to the regulation by the slowly-varying gene expression dynamics.

## Results

Following the earlier theoretical studies on the developmental dynamics to form stripes (Fujimoto et al, 2008), we performed numerical evolution of gene expression dynamical systems (Kohsokabe and Kaneko, 2016). In this article, we review these results and discuss how they support evolution-development congruence as pioneered by Haeckel.

Cells were aligned in a one-dimensional space, each of which consists of a set of proteins. The cellular state was represented by the concentrations of these protein species, whose temporal changes are governed by intra-cellular gene regulation network (GRN) and cell-to-cell interaction via diffusion of some of the expressed proteins. These gene-expression and diffusion processes determine the developmental dynamics to generate a spatial pattern of the gene expression along the aligned cells. From these, fitness was determined from the expression of a prescribed output gene across cells in the one-dimensional space. After each gene expression pattern reached a stationary state through the gene-expression dynamics (i.e., development), the difference between this output expression pattern in space and a predetermined target pattern was computed. If the output pattern matched the predetermined target, the fitness was considered at its highest value; consequently, fitness decreased as the difference became larger.

We identified 100 individuals of virtual organisms with slightly different GRNs and carried out numerical evolution experiments to select those individuals (GRNs) that give rise to higher fitness. From these individuals, the offspring were generated, which had slightly different GRNs mutated from the mother. This was an evolutionary procedure in one generation. From the individuals thus produced, the procedure was repeated to obtain the next generation. (See Figure 2 for simulation procedure. For details, please refer to Kohsokabe and Kaneko, 2016.)

**Figure 2.**
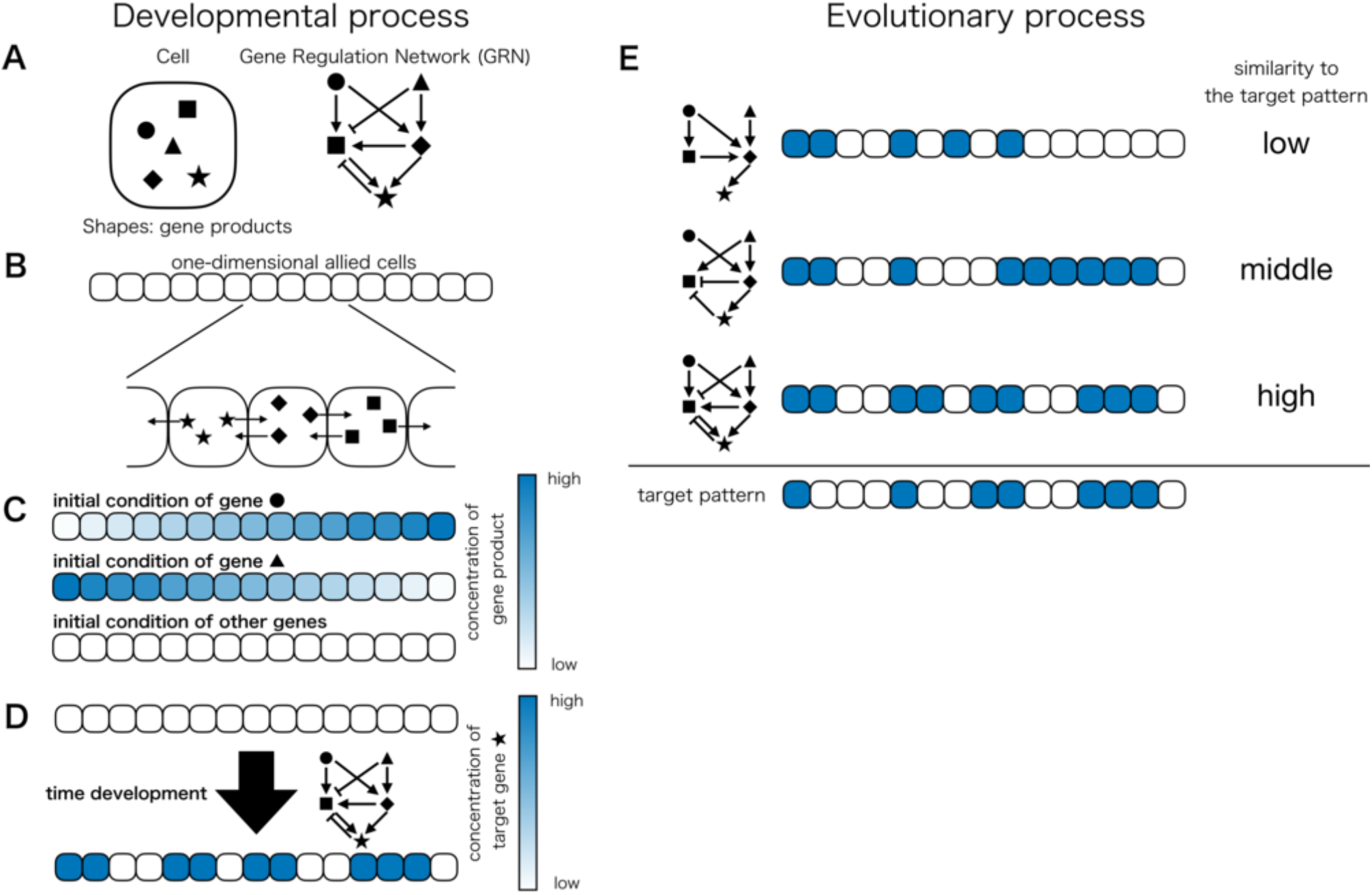
Schematic representation of simulation procedure. Left side: Developmental process (A): Cells contain a variety of proteins, which are coded by genes in the genome. Each shape inside the cell represents a protein that is translated from a different gene. All the proteins in our model are transcription factors that regulate expression of genes encoding other transcription factors. Mutual gene regulation is represented by the gene regulation network as determined by the genome. Gene regulation network consists of edges for activation or inhibition of other genes. (B): An organism consists of cells aligned in a one-dimensional chain. Within an organism, every cell has an identical gene network. Gene products in a cell can diffuse to its neighboring cells. (C): In the initial state (organism in the first phase of development), all genes, except for two, are not expressed. The two genes (represented by a circle and triangle) are expressed to shape the spatial gradient (blue shading), functioning as external morphogen input. (D): The gene expression state of the organism changes through time and space according to the GRN. After sufficient time steps, all the gene expressions reach stationary states. The expression levels are dependent on cells, providing a spatial pattern of expression levels. The phenotype of the model used is given as the spatial pattern of one prescribed output gene (represented by a star; herein, termed output gene). Right side: Evolutionary process (E): Different phenotypes arise from different networks. Fitness is defined by the difference between the output expression pattern and the predetermined target pattern, with the highest fitness values defined by the best match. Such high-fitness individuals can have more offspring. In the next generation, the network edges are slightly altered by mutation.

This numerical experiment can trace how a final target pattern evolved along a single-chain phylogeny from an ancestor to descendant, as in Figure 1. From the highest fitness individual achieved through evolution, we traced back the evolution of individuals to the ancestor to obtain a sequence of output gene expression over generations. We compared this evolutionary sequence with the developmental time-series of the output-gene expression pattern of the fittest individual.

Four examples of such comparisons are shown in Fig. 3, where the space-time diagram of the expression pattern of the output gene is displayed. For development, dynamics of the output-gene expression of the fittest individual is shown with the horizontal axis as the developmental time and the vertical axis as the cellular index (i.e., spatial position), whereas for evolution, the terminal pattern (i.e., after development) of the output-gene for each ancestor through the evolutionary course are displayed, with the horizontal axis as the generation. The similarity between the two space-time diagrams is clear, as they only differ at one stripe or less among all space-time pixels for most cases. Indeed, for 95% of simulation runs, the space-time diagrams between developmental and evolutionary processes showed remarkable similarity. Hence, we found parallelism between evolution and development along the single-chain phylogeny (herein, evolution-development (evo-devo, in short) congruence). Note that this is not identical with Haeckel’s recapitulation theory because this adopts a single-chain-phylogeny comparison, instead of a species-wide comparison.

**Figure 3.**
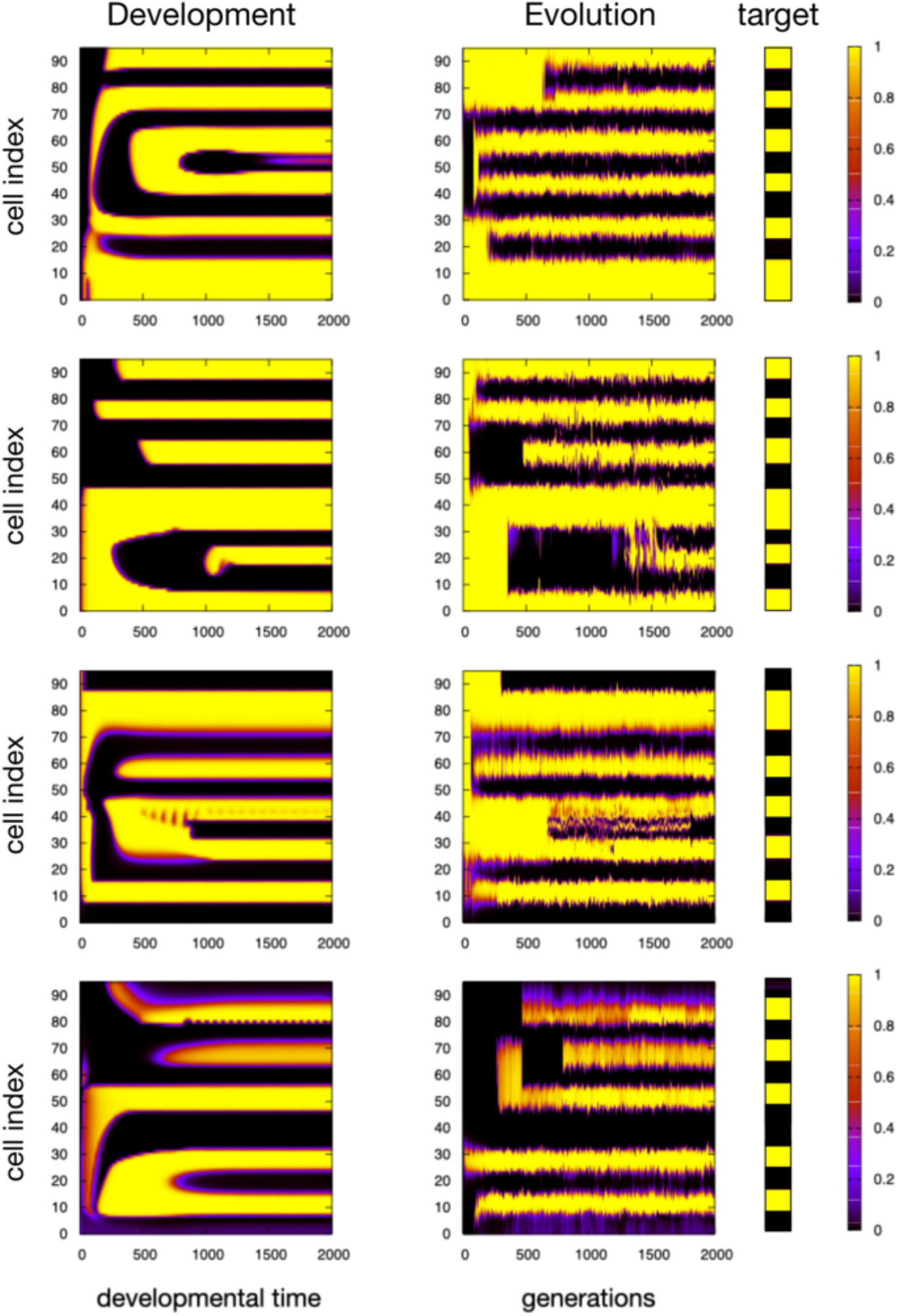
Examples of evolutionary-developmental congruence represented by space-time diagrams derived from our evolution simulation of four different runs. Each row denotes a case. Development: The expression level of the output gene is shown with developmental time as the horizontal axis, and cell index (spatial position) as the vertical axis. The expression level of the output gene of the corresponding cell at a given time is color coded (right sidebar), with black indicating the lowest and yellow indicating the highest expression levels. Development consists of a few epochs with rapid changes in the pattern, separated by quasi-stationary regimes with minor change in the pattern, until the target pattern is shaped by development. Evolution: The spacetime diagram of the evolutionary course corresponding to (A). The expression level of the final output gene (state at developmental time = 2000, for each ancestor) is shown with evolutionary generation as the horizontal axis, and cell index (spatial position) as the vertical axis. This figure shows how the terminal pattern after development changes through evolution. At each generation, the final pattern of the direct ancestor is shown (See also Fig.1). The evolution of the developed output pattern consists of quasi-stationary regimes sandwiched by epochs with rapid change resulting from mutation until the target pattern is evolved.

### Development with epochs that corresponded to those derived through evolution

Here, we discuss the correspondence between developmental and evolutionary space-time diagrams in our simulations.

Pattern formation remarkably progressed in a stepwise manner with respect to evolutionary generation and developmental time. Each stripe emerged discretely rather than gradually. Except for these epochs, pattern changes were rather small and the pattern remains quasi-stationary between epochs.

The formation of epochs in evolutionary courses is reasonable. Because genetic mutation causes change in GRN, a change in the reaction also occurs (the rule of dynamical systems). In the present model with strongly purifying selection, only the neutral or beneficial mutations remain during evolution. However, beneficial mutations are rather rare, and many generations are required for them to occur. Furthermore, the accumulation of neutral mutations is often needed for beneficial subnetwork to formulate. Once such relevant mutations occur, the pattern can make a drastic change. Thus, the evolutionary course of the developed pattern consists of a quasi-stationary regime and requires several epochs to change the stripe pattern. This epochal pattern change in evolution has previously been coined as “punctuated equilibrium” (Eldredge and Gould, 1972).

In contrast, in development, there is no a priori reason why the process consists of a long quasi-stationary regime and several epochs with drastic changes. Here, we have found that after evolution, gene(s) whose expression much more slowly changes emerges, affecting the expression of the output gene. Here we will term such gene as “slow gene”. During the quasi-stationary regime, the output can change only slowly due to this slow gene(s), whereas a small change in the expression of this slow gene(s) brings a drastic change. This is due to bifurcation in dynamical systems (as will be explained below). Such gene(s) with slowly varying expression always emerge as a result of evolution, and it functions as a bifurcation parameter to control the fast changes in the expression of the output gene.

### Bifurcation behind evo-devo congruence

In dynamical systems, drastic, qualitative change in attractors (final states) induced by slight changes in control parameter(s) is referred to as bifurcation (see for instance, Hirsch, Devaney and Smale, 1974; Strogatz, 1994). For most parameter regimes, the attractor continuously changes as parameters change but without qualitative change. However, when a parameter reaches a certain value, a bifurcation, qualitative change in the attractor occurs, e.g., change from one type of fixed-point attractor to another, or from fixed-point to a limit-cycle attractor with oscillatory expression.

The correspondence between evolution and development is explained in terms of bifurcation (see Fig. 4 for the bifurcation observed in our model). In development, slow change in expression that controls downstream genes works as a bifurcation parameter, and bifurcation in the expression dynamics generates a novel steady state, which gives rise to an epoch. In evolution, when a relevant mutation occurs, qualitative change in the final state occurs as bifurcation, and this novel state generates an epoch. Thus, both the epochs in development and evolution are generated by the same bifurcation. In this way, evo-devo congruence is explained by a common bifurcation at each epoch. Indeed, in most examples we have examined, the same bifurcation occurs to generate a novel stripe for both development and evolution, as in Fig. 4.

**Figure 4.**
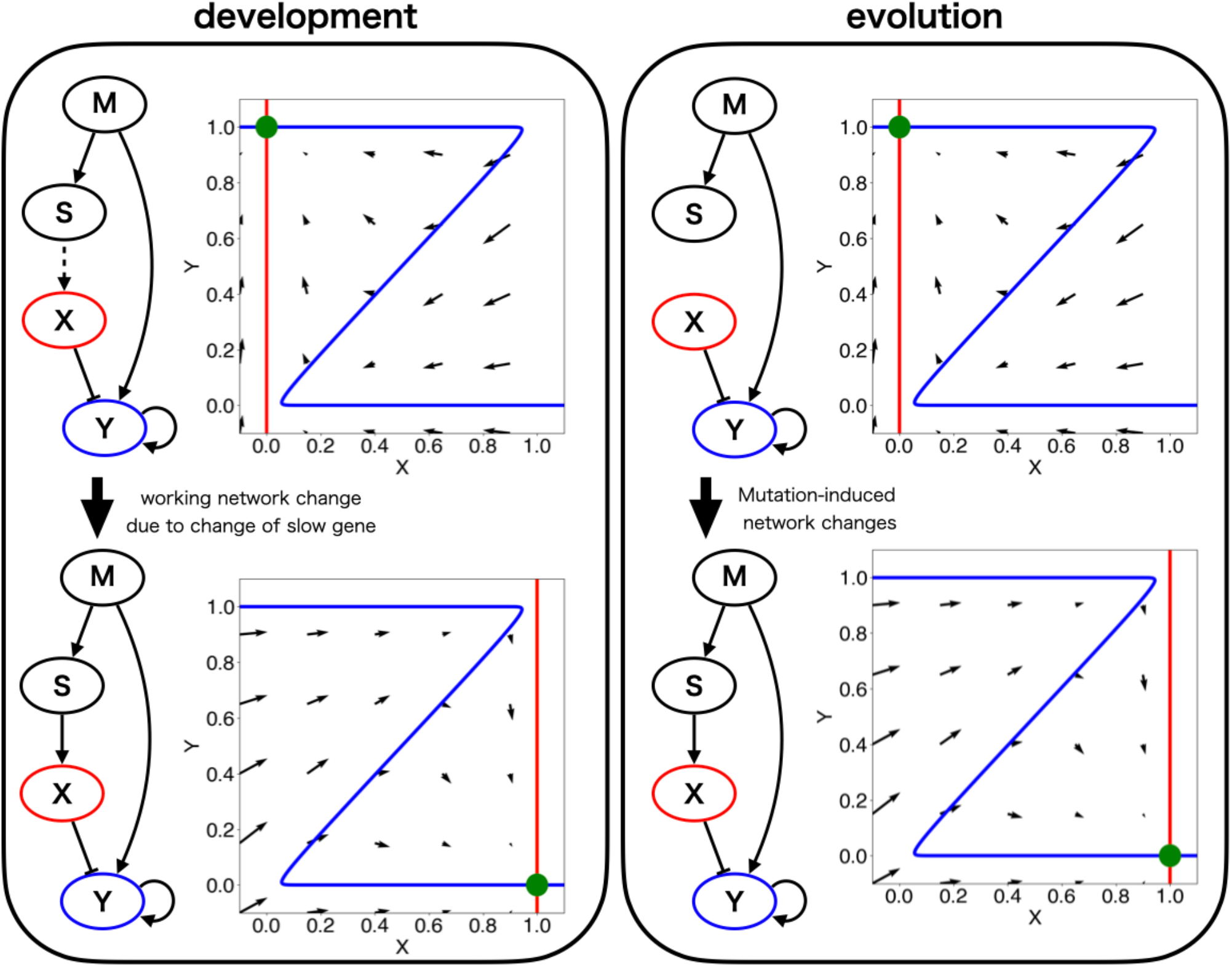
Bifurcation of fixed points during development (left) and evolution (right). In the left side, the network structure that is responsible to the expression of the output genes at the moment is displayed. Changes in the flow of the expression levels of two genes X and Y in the shown GRN are plotted. The horizontal axis shows the expression level of X, whereas the vertical axis shows that of Y. The red line represents the nullcline of gene X (i.e., the temporal change in the expression level of gene X vanishes), whereas the blue line represents the nullcline of gene Y. The green circle at the intersection of the nullclines is the fixed-point attractor, the final stable cell state. Left: Developmental change. The upper phase diagram represents an earlier stage before an epoch, with the GRN on the left. The expression of gene S is lower than the threshold for the expression of gene X (as indicated by the dotted regulative arrow from gene S to gene X on the left network), the stable fixed point, as given by the intersection of nullclines at approximately (0, 1). As development progresses, the expression level of gene S increases. Once its expression exceeds the threshold of gene X, the nullcline of X shifts slightly to the right, indicating a higher expression level. The flow diagram at this later developmental stage is plotted at the bottom, with the responsible GRN on the left. GRN that works effectively alters according to the change in the expression level of the slow gene (with the solid arrow from gene S to gene X). Gene X inhibits the expression of gene Y, and the fixed point changes to (1, 0). Now the fixed-point attractor has shifted from upper left to lower right because of bifurcation, resulting from the change in the slow expression level that works as a bifurcation parameter. Evolution (right): The phase diagrams for the expression levels of genes X and Y with the corresponding nullclines are displayed before and after the relevant mutation in the evolutionary course. In the upper column, the path from S to X had not yet been acquired in the GRN. Hence, the expression level of gene X is not activated. As shown in the corresponding phase diagram, the stable fixed point lies approximately at (0, 1). In the later generations, after the relevant mutation occurs, the GRN has acquired the positive regulative path from gene M to gene X. Because the input to X is now sufficiently large, the phase diagram has changed (bottom). The expression level of gene X is higher, so the fixed point moves to (1, 0). By comparing the right and left diagrams, a strong correspondence is discernible between both bifurcations through evolution and development, as well as the corresponding change in gene networks.

### Expansion of working networks

Gene expression dynamics are driven by the gene regulation network (GRN), in which each protein expression mutually activates or inhibits. In GRN, the generation of spatial patterns has two classic mechanisms, feed-forward and feedback regulations (see Fig. 5).

**Figure 5.**
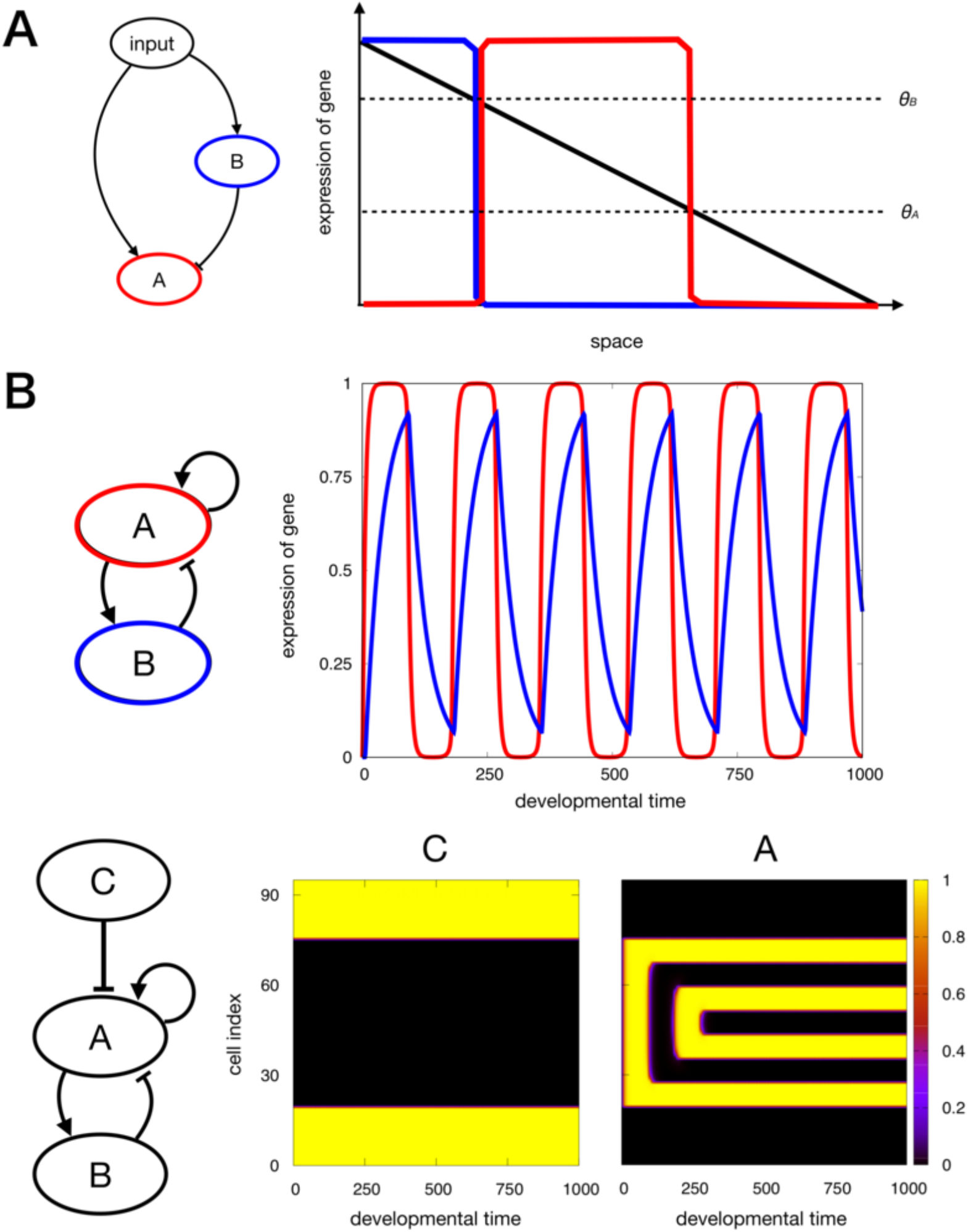
Feedforward and feedback networks. A. Left: An example of incoherent feedforward network, which reads the external morphogen gradient as an input and forms a stripe. Right: Spatial expression of each gene due to the regulation of the left network. The red line corresponds to the expression level of gene A, whereas the blue line corresponds to that of gene B. θ_A_ and θ_B_ are the expression threshold for each gene. B. (top) Minimal network for the oscillatory expression with the time series of the expression for a specific cell. Gene A activates the expression of gene B and itself, whereas gene B suppresses gene A. Without interaction with other cells (i.e., no effect of spatial boundary), the expression level oscillates in (developmental) time as plotted in the right column, with developmental time as the abscissa and the expression levels of A (red) and B (blue) as the ordinate. (bottom) Spatial fixation of the oscillatory expression under a fixed boundary: The input from gene C, which was influenced by a maternal factor, is included in the oscillatory network. The space-time diagram of genes C and A illustrate how the oscillatory expression of gene A is fixed to form spatial stripes, inside the region whose boundaries are settled by expression of gene C.

The classic mechanism for stripe formation, the feedforward regulation, was analyzed in the segmentation process in Drosophila (von Dassow et al., 2000; Jaeger et al., 2004; Ishihara, Fujimoto and Shibata, 2005). Here, a gene receives input from the morphogen gradient as spatial information, to induce an “on/off” response under a given threshold level, so that the gene is expressed on the one side of space, and non-expressed on the other side. Then, another “downstream” gene receives positive (or negative) input from this gene and negative (or positive) input from the morphogen, responding to create another segmentation in space (see Fig. 5a). Combining these feedforward regulations results in formation of more stripes for the downstream genes.

The other mechanism for stripe formation commonly observed in our simulations takes advantage of feedback regulation. In a negative-feedback regulation (see Fig. 5b), expression level can exhibit temporal oscillation. In a system with spatially arranged cells with mutual diffusion, temporal oscillation at a single-cell level is fixed into a spatial periodic pattern under an appropriate fixed boundary condition (Kohsokabe and Kaneko, 2017). Note that though they may look similar, this dynamics is different from so-called “clock and wavefront” model (Cooke and Zeeman, 1976); unlike clock and wavefront model, this mechanism does not require a moving global morphogen to propagate the spatial pattern. Only local cell-to-cell diffusion is used. This mechanism is triggered by static local morphogen at the boundary of the region where the pattern is formed.

The stripe formation processes evolved in our model could be generated by sequentially combining the two mechanisms. For the feedforward network to work, spatial gradient in the upstream network is necessary; for the feedback mechanism to work, the fixed expression at a boundary for a certain domain has to be established in advance to fix the temporal oscillations to spatial stripes. To produce the boundary, the upstream feedforward mechanism is needed to read the external morphogen gradient.

Through evolution, GRN is successively expanded downstream of the input and added to generate further stripes. This works as long as the upstream mechanism is not affected by the added downstream network. In the evolutionary course, this can be achieved by the successive addition of downstream networks.

In the development of the fitted individual, all the components in the GRN exist at the beginning. However, if the downstream network is only activated later by a “slow gene” and not at the start of development, the developmental process can proceed similarly as the evolutionary process.

To understand the role of genes with slowly varying expression, a core part of the GRN responsible for stripe formation at each epoch, was extracted. The core network at each epoch, termed as the “working network,” is successively activated by the slowly changing gene expression (see Fig. 6). Indeed, the “slow gene” that influences the target gene serves as a control variable for the switch in output gene expression dynamics. As its expression level changes slowly, some downstream genes are activated (or repressed), so that working networks expand in the same process as evolution. Hence, the ordering of working networks over epochs is consistent with development and evolution, which progress from a feedforward-based network to network including feedback loops. The validity of this ordering has been confirmed statistically (see Kohsokabe and Kaneko, 2016 for detailed analyses).

**Figure 6.**
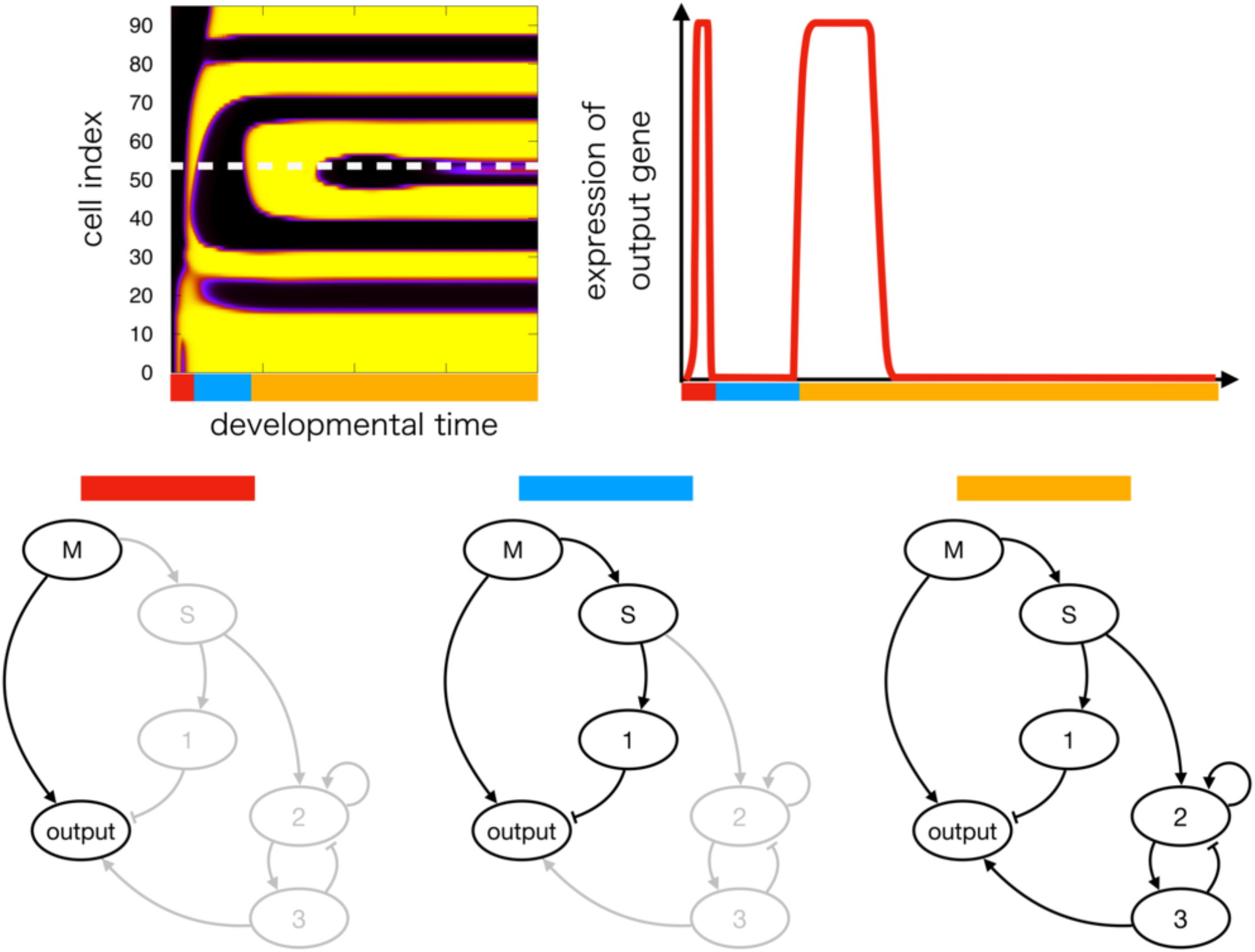
Schematic representation of the expansion of working networks. With the slow change in the expression level of a certain gene that influences downstream genes, the working network expands. Upper left: An example of space-time diagram of the development of an individual that reached the highest fitness after evolution. Here, we focus on the dynamics of the gene expression of one specific cell (white dashed line). The expression of the output gene of the cell is displayed in the right. The stepwise change in the expression is discernible and thus, epochs in development are detected. Three temporal regimes separated by the epochs are represented as red, blue and orange bars plotted against time. Upper right: The dynamics of the output gene are shown, with the horizontal axis as developmental time and the vertical axis showing the expression level of the output gene. Bottom: The working network that influences the output expression at each time regime is indicated by its corresponding colored bar.

The “expansion of working networks” is consistent with the bifurcation we discussed in the previous section. Through the slow change in upstream gene, the expression dynamics at the downstream network drastically change via bifurcation, leading to novel stripe formation.

### Slowly changing gene expressions

We have already mentioned the relevance of gene(s) whose expression changes slowly and controls the downstream expression of genes to achieve evolution-development congruence. We examined 500 examples and have found that the networks after evolution always include a gene with slowly changing expression that is essential to produce an epoch. The time scale of this expression is slowing down during evolution (see Fig. 7). The gene with slowly changing expression did not give a direct input to the output gene, but instead, gave an input to a gene that provides input to the output gene.

**Figure 7.**
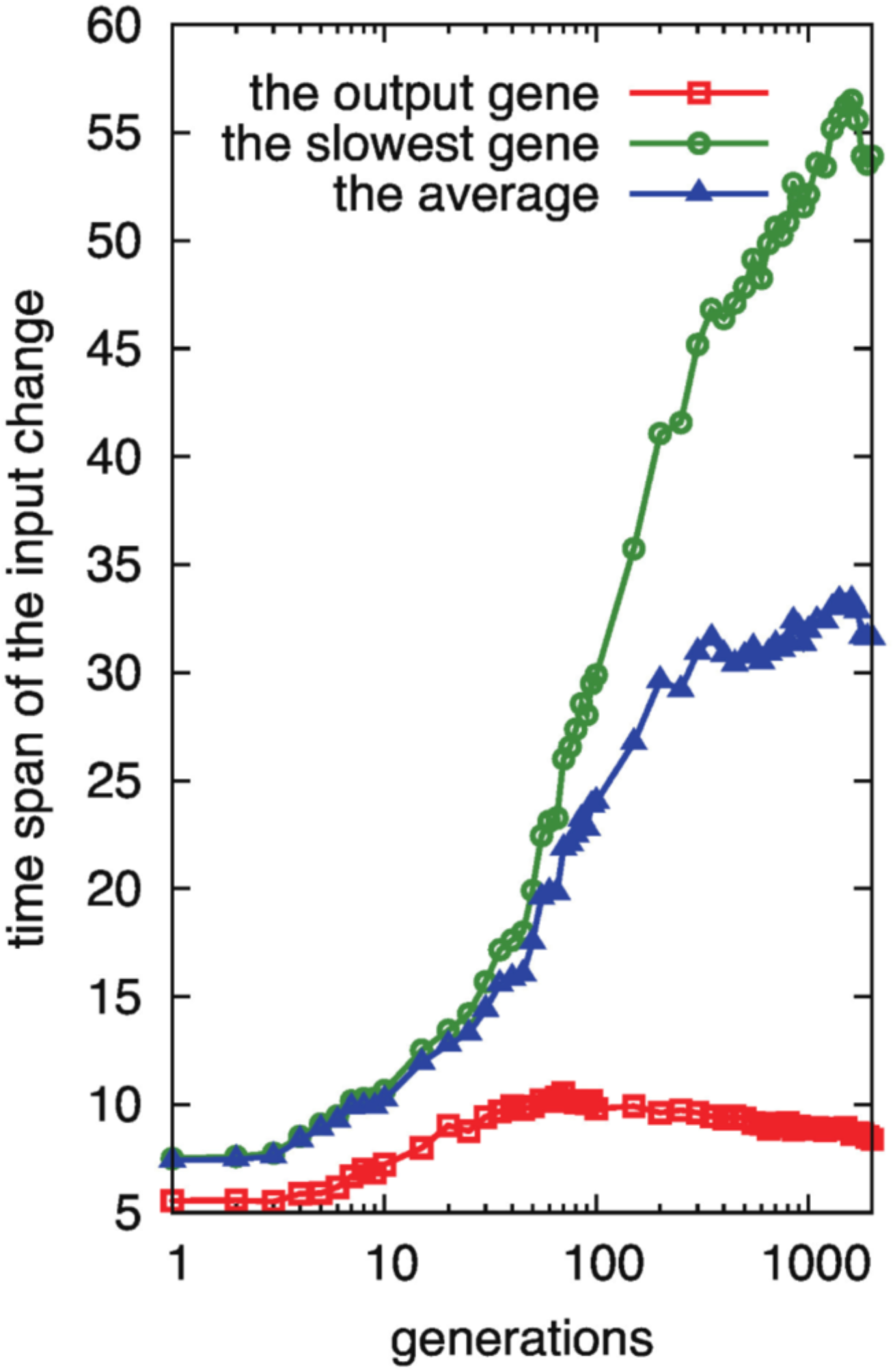
Evolution of the time scale for the input for the output and other genes. Using the genes whose expression levels change between on and off for each developmental epoch, the time scales are computed as the time span that the input for the gene passes through the dynamic range during each development. The red square gives the time span for the input of the output gene, whereas the blue triangle (green circle) denotes the average (the largest) time span for input among the genes that have a path to the output gene. The time spans are computed from an average of 500 runs of evolution simulations. (reproduced from Kohsokabe and Kaneko, JEZB 2016)

Further, we statistically confirmed the evolution of these genes with slowly changing expression that controls the output gene. In the present model, such slow changes evolve through a decrease in the rate constant of a certain gene expression or through a change in the expression threshold so that the expression level stays close to it. Besides this slow change within a cellat a single-cell level, the slow expression change can be propagated in the entire space through diffusion to other cells to bring the bifurcation of the pattern.

### Heterochrony induced by the alteration of the timescale of slow genes

As mentioned above, after evolution, developmental change of expression of some genes always take much slower time scales than other genes. The expression dynamics of such ‘slow’ genes dominantly control the developmental timetable. Mutational changes occurring in the downstream regions of these slow genes may often be beneficial or at least neutral, as the effects of such mutations are not apparent until later developmental stages, so that the already-acquired pattern would be conserved.

Then, what will happen when the timescale of such slow genes is altered by further evolution? Timescales of genes could be manipulated by changing the parameters for each gene in our model. An example of such manipulation is shown in Fig.8; the original developmental dynamics is given (Fig.8A). In this example, one slow gene controls the whole timetable of the development, especially the timing of the last developmental process to make the last stripe (down regulation of the target gene in the middle of the space). The developmental dynamics by shortening the timescale of the slow gene (i.e., to make faster change in gene expression) is shown in Fig.8B, whereas that by lengthening the time scale (i.e., to make slower change in gene expression) is given in Fig.8C and 8D. It is discernible that the last developmental process is scheduled earlier (B) or later (C and D) than the original case as the timescale of the slow gene is manipulated. For (B) and (C), the developmental process, i.e., the stripes and ordering of their formation, is preserved, and only the speed of the development is altered. In contrast, in the case D, where the timescale parameter of the slow gene is changed to a much larger value, the last developmental process would no longer occur (or the duration of the second last process would be infinite). In other words, the terminal pattern in the case D is similar to that of the earlier stage of the original one, hence the ‘juvenile’ pattern of the original is frozen. This corresponds to the neoteny, a typical example of heterochrony, and an important driver of the evolution of development (Gould, 1977; Hall, 1999).

**Figure 8.**
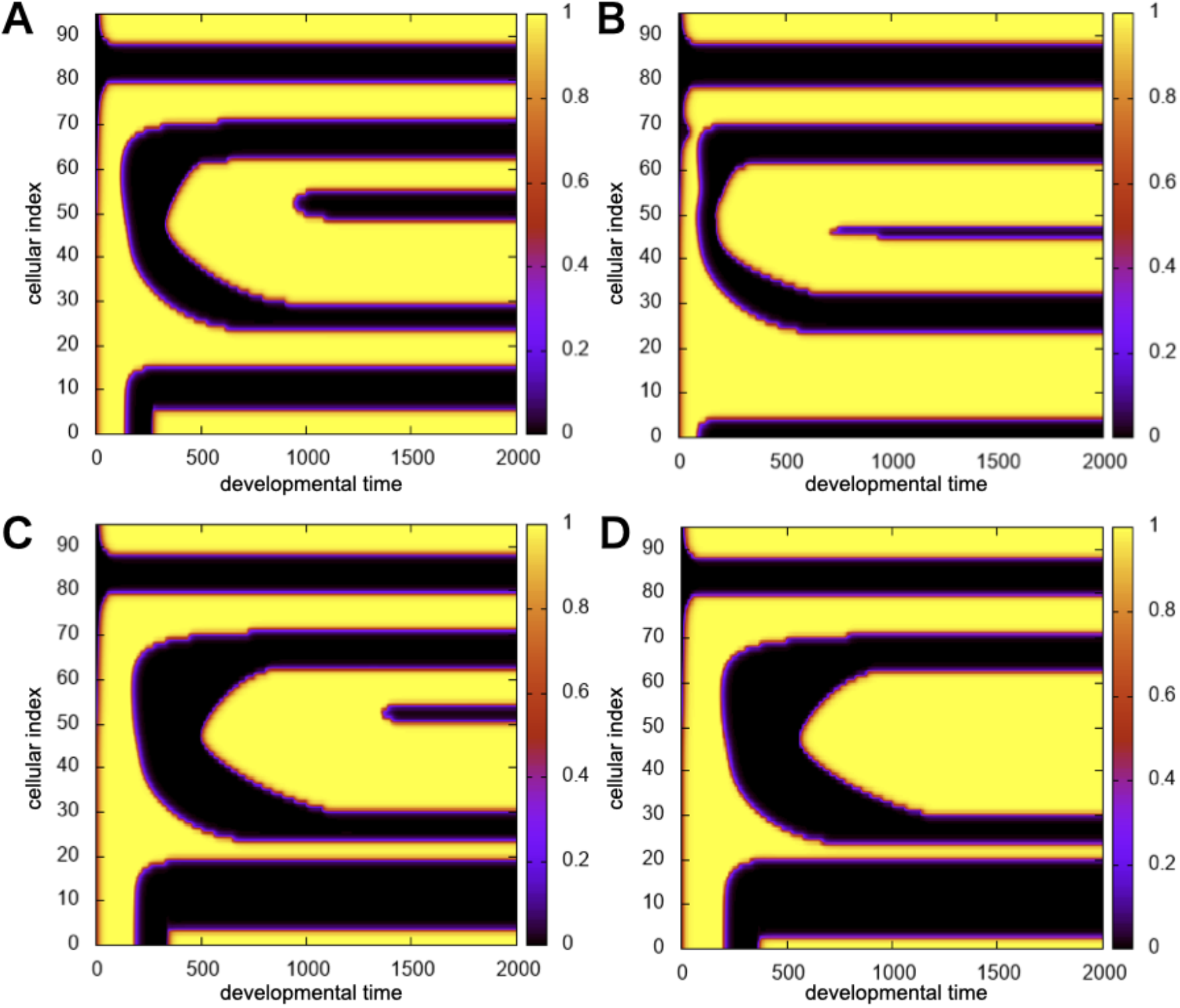
Heterochrony induced by the alteration of the timescale of the slow gene. (A). Space-time diagram of the target gene expression with developmental time (horizontal axis) and cellular index (vertical axis). The color represents the expression level of the target gene, given by the color scale at the right bar, as in Figure 3. The timescale parameter of the slow gene for the expression dynamics is set at 10. (B)-(D). Space-time diagram of the target gene expression with the same network as in (A) but with a different timescale of the slow gene. Indication by axes and the side color scale are the same as in (A). (B). The time scale parameter of the slow gene is set at 5, smaller than in the case (A) (C). The time scale parameter of the slow gene is set at 25, larger than in the case (A). (D). The time scale parameter of the slow gene is set at 28.5, larger than in the case (C).

In general, as long as the change in the timescale is moderate, only the duration of some developmental processes is shortened or prolonged; the overall developmental course and final pattern are preserved. Then, the skipping of the last process could occur, should the timescale of the slow gene be further extended. Of course, if the timescale is changed too much, the original developmental course could be destroyed at some point.

To sum up, the evolution-development simulations in this study suggest that behind the heterochrony in evolution, slow genes that evolved to control the timetable of developmental process are shared among some clades.

### Violation of evo-devo congruence

Although evo-devo congruence was frequently observed and has been discussed in terms of the expansion of working networks and of the correspondence of bifurcations of gene-expression states in terms of bifurcations of gene expression state(Fig.4) and mechanisms of pattern formation(Fig.5) as well as expansion of working networks by slow control gene(Fig.6), a few cases (approximately 5%) deviated from this evo-devo congruence, as shown in Fig. 9. In this case, during the developmental process, the first and fourth upper stripes were branched early, and the second and third stripes were branched from the boundary generated by the first and fourth stripes. These stripes appeared independently during evolution (see also Kohsokabe and Kaneko, 2016 for another example).

**Figure 9.**
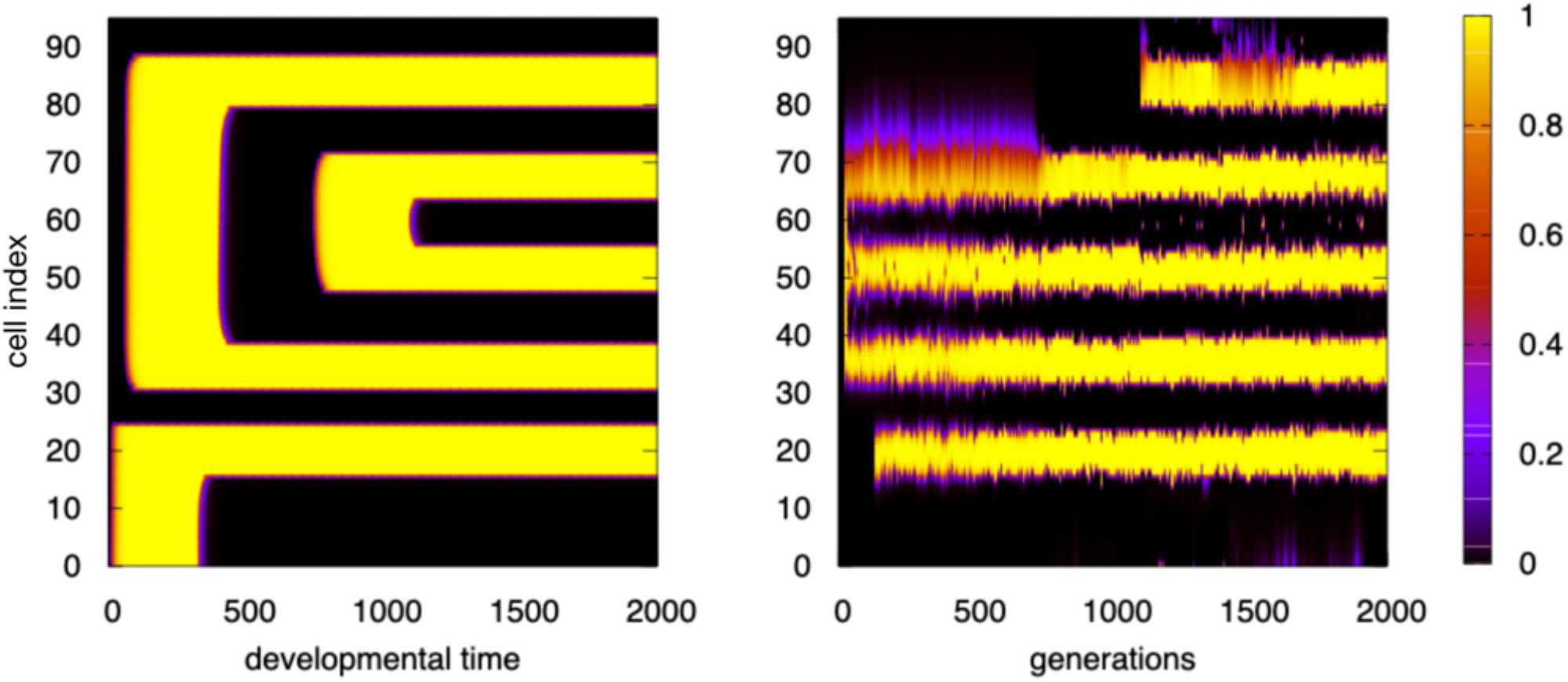
Violation of evo-devo congruence. Development: Space-time diagram of the expression with developmental time (horizontal axis) and cell index (vertical axis) and the color scale (right bar, as in Figure 3). The first and the fourth upper stripes emerge together during the early stage in development. Following this, the second and the third upper stripes are formed concurrently. Evolution: The expression level of the final output gene is shown with generation (horizontal axis) and cell index (vertical axis) and the color scale (right bar, as in Figure 3). The three middle stripes are formed concurrently at the earlier stage of evolution. Later, the top and bottom stripes are formed independently of these three stripes. By comparing the right and left diagrams, evo-devo congruence is topologically violated.

The violation of the congruence was due to a mutation in an upstream expression, causing a change in the boundary condition for the feedback oscillation of the downstream expression gene. (In the example above, the upper four stripes were generated from the fixation to spatial pattern of a temporal oscillatory expression). We have studied several other examples deviating from the evo-devo congruence and have confirmed that the differences in the order of emergence and/or branchings of stripes between development and evolution is caused by an upstream mutation that altered the boundary condition for the feedback mechanism.

## Discussion

From the evolutionary simulations and dynamical systems theory, we have demonstrated evodevo congruence using single-chain phylogeny. Theoretically, the comparison between development and evolution using this phylogeny approach is straightforward; however, caution against species-wide comparison, which Haeckel adopted for the recapitulation, is essential. Here, if the evolution and development progress in a stepwise manner by expansion of working networks, as governed by common bifurcation in the expression state, then, the species stemming from each branch in the phylogenetic tree (in Fig.1) are expected to retain the ancestor’s epochs during development as long as bifurcation is preserved. Evo-devo congruence in the single-chain phylogeny will then be mapped to the parallelism of evolution and development across species, i.e., those evolutionarily diverted from a common ancestor. This may provide a possible explanation of what Haeckel had reported.

At a first glance, our results, which may support Haeckel’s theory, does not fit the standard viewpoint in the evolutionary developmental biology, especially because Haeckel’s recapitulation theory has long been dismissed or has been regarded as a historical mistake. However, history proves that Haeckel’s theory was not rejected, but rather test for its authenticity had been forgotten as it has been rendered obsolete in comparison to experimental embryology as pioneered by Roux and with links to molecular biology(Gould, 1977). During Haeckel’s time, the mechanism of heredity was unknown and there were no distinctions between genotype and phenotype. Hence, elucidating how genes control development was not possible, and Haeckel’s theory could not be proven or disproven. Fortunately, we now have access to experimental data and theoretical analyses to investigate his hypothesis quantitatively, and more importantly, scientifically.

At present, we can now verify evo-devo congruence by investigating the stepwise epochal changes with bifurcation in the developmental process. First, the quantification of changes in gene expression patterns during development is needed and can be achieved by using transcriptome analysis. One can then examine whether the changes occurred in a stepwise manner across several epochs. Next, the gene regulation network (GRN) for each species should be explored. Although its complete extraction might be difficult, it can be partially estimated and the expansion of working networks can be studied. Lastly, by analyzing species stemming from common ancestors, the validity of evolution-development congruence can be determined. As for the bifurcation, although dynamical systems analysis per se may be difficult, it can be estimated using the gene expression changes at each epoch and check if these changes occur in a stepwise manner.

Because we can only observe the development and morphology of the present organisms that diverged from common ancestral species, developmental dynamics cannot be easily traced through evolution (See Fig. 1). Thus, the correspondence between evolution and development through common bifurcation is not so easy to verify from experimental data. However, if the morphological novelty was a result of bifurcation, then different novel morphologies are expected to diverge from a common ancestral pattern. This is consistent with Haeckel’s theory or with von Baer’s third law of embryology, which claims that a common basic morphological feature emerges prior to special features for each species.

In our approach, developmental change is analyzed only in terms of gene expression patterns, i.e., intra-cellular chemical concentrations. However, the motions and arrangement of cells to form tissues are essential in the development to an adult form. Indeed, extensive studies on evolution-development correlation have been carried out using detailed anatomical comparison (Willmer et al., 2009). Although changes in gene expression patterns may underlie the spatial configurations of cells, this remains to be a limitation in our analysis. However, even though the models discussed here are concerned with gene expression dynamics, analysis based on dynamical systems is rather general. Pattern formation, including cell arrangement and tissue organization, is also represented by dynamical systems, wherein the state is not restricted to gene expression(Murray, 2002). Even if such cellular dynamics are included, evolution is represented by successive changes in dynamical systems to generate patterns with higher fitness. Thus, stepwise changes in morphological dynamics are expected. Based on the emergence of slow variables and bifurcation, our theory can thus be generally applied to any morphological dynamical system. Hence, the demonstrated evo-devo congruence will be valid beyond the gene-expression models. The search for a stepwise developmental process and a control of slow variables in general morphological processes will deepen our understanding of the evolution-development relationship.

Slow process during development is essential for evolution-development congruence. Interestingly, in several evolution models, including catalytic reaction, gene expression, and pattern formation dynamics, the existence of one or few slow modes, which regulate the other processes, is also observed (Furusawa & Kaneko, 2018; Sato and Kaneko, 2020). In fact, if all modes occur at the same time scale, fitting complex developmental dynamics to direct to a certain pattern would be difficult, similarly to the proverb ‘too many cooks spoil the broth’. If slow modes affect many other processes relevant to development, it will be easier for the directed process to reach a complex pattern. In addition, the existence of slow modes may have facilitated evolution since changing the mode would affect many other processes. Since evolution occurs at a much slower time scale than development, the slow mode in development provides the fastest direction in evolution, so that evolution and development can be connected. We expect that the exploration of slow processes in developmental data and their comparison across species will be important. This can be facilitated by transcriptome analysis and multicellular morphological processes.

As an example of the observed developmental process that suggests the regulation by slowly changing gene expression, we pick arthropod segmentation. As described in the earlier section, the modeling that combines the gene expression and pattern formation dynamics was originally designed for theoretical studies of segmentation (Salazar-Ciudad et al., 2001b; François, Hakim and Siggia, 2007; Fujimoto et al., 2008). In particular, segmentation of Drosophila melanogaster, was analyzed by fitting the parameters in gene expression dynamics, via the use of experimental data (Perkins et al., 2006; Manu et al, 2009; Gursky et al., 2011). In contrast, the evolution of gene regulation networks we discussed here assumed neither a known network structure nor the parameters in the observed reaction dynamics. However, we can see commonality between the results in our abstract model and the segmentation of Drosophila melanogaster.

It is now considered that the initial expression of hunchback determines the afterward gap gene expression state during the segmentation of Drosophila melanogaster. Of note, the hunchback expression decays slowly in time, affecting the transient expression dynamics of other gap genes. Verd et al. (2018) introduced a dynamical systems model based on the experimental data, where the expression dynamics of gap genes (Krüppel, giant, and knirps) depend on the continuous change in the initial value of the hunchback expression. Importantl’sy, they show qualitative changes as the hunchback expression is changed. This is consistent with the results we described in the present paper; the hunchback expression controls the bifurcation as a slow gene.

Therefore, the evolution of segmentation patterning of flies could be achieved by changing parameters that regulate the genes downstream of the hunchback, which would generate epochal, rather than continuous changes in phenotype, as in the bifurcation. In fact, in an evolution simulation by Rothschild et al., (2016), changes in parameters regulating the expression of even skipped, which is located downstream of hunchback, often bring the first epochal phenotypic change toward further segmental pattern changes.

The theoretical studies we reviewed here also have implications to the developmental process. With the slow change in hunchback expression, the gene-regulation network that works at each moment will be expanded in a stepwise manner, from the structures that consist of only feedforward interactions to those containing feedback loops. It will be interesting to examine this prediction via transcriptome and network analyses through the segmentation process.

Lastly, the developmental hourglass, a bottleneck in development, has received attention recently, where the phenotypic differences across species (stemming from the common ancestor) are minimal (Irie and Kuratani, 2014). In the embryo, the difference is larger and decreases during development up to the phylotypic stage, and later the difference increases. Because we have set a unique target pattern as the highest fitness state and have carried out strongly purifying selection, our numerical evolution model cannot be used to verify the hourglass hypothesis. However, by relaxing the fitness conditions to allow for phenotype diversification, the hypothesis can be tested, and we have already collected preliminary data that support the developmental hourglass model. Particularly, in the later developmental stage from the bottleneck, we have found the evolution-development congruence. Accordingly, Haeckel’s theory does not necessarily contradict the hourglass model.

## Acknowledgments

The authors would like to thank Shigeru Kuratani, Naoki Irie, Masahiro Uesaka, Tetsuhiro Hatakeyama, and Shuji Ishihara for the stimulus discussion. This work was partially supported by the Grant-in-Aid for Scientific Research on Innovative Areas (grant number: 17H06386) from the Ministry of Education, Culture, Sports, Science and Technology (MEXT) of Japan.

